# Polaris: a universal tool for chromatin loop annotation in bulk and single-cell Hi-C data

**DOI:** 10.1101/2024.12.24.630215

**Authors:** Yusen Hou, Audrey Baguette, Mathieu Blanchette, Yanlin Zhang

## Abstract

Annotating chromatin loops is essential for understanding the 3D genome’s role in gene regulation, but current methods struggle with low coverage, particularly in single-cell datasets. Chromatin loops are kilo-to mega-range structures that exhibit broader features, such as co-occurring loops, stripes, and domain boundaries along axial directions of Hi-C contact maps. However, existing tools primarily focus on detecting localized, highly-concentrated, interactions. Furthermore, the wide variety of available chromatin conformation datasets is rarely utilized in developing effective loop callers. Here, we present Polaris, a universal tool that integrates axial attention with a U-shaped backbone to accurately detect loops across different 3D genome assays. By leveraging extensive Hi-C contact maps in a pretrain-finetune paradigm, Polaris achieves consistent performance across various datasets. We compare Polaris against existing tools in loop annotation from both bulk and single-cell data and find that Polaris outperforms other programs across different cell types, species, sequencing depths, and assays.

## Introduction

The three-dimensional genome conformation is vital for cellular processes, including gene expression and DNA replication [1, 2, 3, 4]. However, its formation, functional implications, and cellular heterogeneity remain poorly understood. Chromosome conformation capture assays [5, 6, 7, 8, 9, 10, 11, 12, 13] have enabled the study of pairwise interactions between DNA fragments in either cell populations or individual cells. They hold great promise for advancing our understanding of 3D genome organizations. Notably, emerging single-cell techniques [7, 8, 9, 10, 11, 12, 13] offer significant potential to uncover cellular heterogeneity. Researchers have observed multi-scale genome organizations, including chromatin loops [14], topologically associating domains (TADs) [15, 16], compartments [5, 17], and architectural stripes [18], using these assays. However, data produced by many of them, particularly single-cell assays, are often too sparse to be effectively analyzed by existing tools [19, 20]. Among these elements, chromatin loops have attracted significant attention [14, 21, 22, 23]. They not only facilitate long-range interactions [24], but also play a crucial role in maintaining the dynamic chromosomal conformations [1, 21].

Many loop callers have been developed [14, 25, 26, 27, 28, 29, 30, 31, 32, 33, 34, 35], evolving from early enrichment-based methods to recent pattern recognition and data-driven approaches, but most of which are designed for bulk data. Enrichment-based methods, such as HiCCUPS [14], identify loops by comparing contact pairs with their local surroundings. While effective in certain scenarios, they lack robustness to noise and often misidentify isolated high-intensity interactions as loops. Chromatin loops frequently appear as blob-shaped patterns in contact maps, which has motivated the development of pattern recognition approaches [27, 28]. Mustache [27] employs multiscale difference of Gaussians, while Chromosight [28] uses a template-matching approach to detect such patterns. Although these methods perform well on typical-coverage Hi-C data, they struggle to identify loops that deviate from blob shapes. To overcome these limitations, supervised learning methods, such as Peakachu [29], have recently emerged, which learn loop patterns directly from labeled data. They usually perform well in detecting loops from bulk contact maps but encounter challenges with single-cell data, where loop signals are substantially diminished.

An effective strategy for analyzing sparse Hi-C data, particularly scHi-C data, is to enhance contact maps. For instance, SnapHiC [31] performs random walk to impute missing interactions, thereby improving loop annotation from scHi-C data, while Higashi [36] models scHi-C data as a hypergraph and imputes missing interactions by predicting missing edges. Nevertheless, loop annotations from these enhanced data remain less accurate compared to annotations derived from bulk data. Other approaches address insufficient sequencing depth by incorporating additional data. RefHiC [33], for instance, leverages a panel of reference data to augment input. While tools that incorporate additional data often achieve high accuracy, their applications are limited by the availability of such supplementary data. Data-driven approaches are often more accurate but can struggle with computational efficiency. To mitigate this, YOLOOP [34] was recently introduced. It significantly accelerates loop annotation by analyzing larger contact map patches. However, this speed comes at the cost of accuracy, with predictions deviating by up to 10 bins from the exact loop positions. Most loop callers rely on local features within small windows to identify loops. While effective in high-coverage datasets, their performance declines in low-coverage or single-cell data.

Here, we introduce Polaris, a universal loop caller that processes large chromosomal contact matrices, naturally integrating both local and global patterns through its model design. Polaris leverages broader features within contact maps, such as loops positioned at TAD corners, co-occurring loops along rows or columns, and overlaps with architectural stripes, allowing it to capture patterns missed by existing tools. This design enhances Polaris’s robustness to noise and sparsity, making it particularly effective for analyzing single-cell and low-coverage datasets. Moreover, Polaris’s novel training paradigm ensures accurate loop annotation across diverse datasets, addressing challenges related to resolution variability, sequencing depth, species differences, and experimental protocols.

## Results

### Polaris is a universal loop caller

Polaris takes a contact map as input and makes chromatin loop annotations. It comprises two components (Fig. 1a, b, Supplementary Fig. 1, Methods): (i) a neural network that assigns loop scores to all pixels of the input contact map, and (ii) a density-based clustering module that identifies discrete loops from these scores. Unlike existing tools [33, 29], Polaris operates on a much larger region (i.e, 224 × 224) and simultaneously predicts loop scores for all pixels within that region, greatly enhancing computational efficiency. The network architecture combines axial attentions [37] and a U-Net backbone [38] (Fig. 1a), providing a large receptive field and focusing on axial interactions for more accurate predictions. We designed the axial attention (Fig. 1b) to capture axial features, such as the co-occurrence of stripes and loops, and observed that it performs as intended (Supplementary Note 1, Supplementary Figs. 2-3). To leverage the abundance of unlabeled Hi-C data, we developed a knowledge distillation-based [39] pre-training and fine-tuning framework (Fig. 1c). Specifically, Polaris was pre-trained on massive Hi-C datasets (Supplementary Table 1) to align with RefHiC, by using RefHiC’s predictions as targets. Fine-tuning was then conducted using downsampled Hi-C data from the GM12878 cell line. Unlike RefHiC, which requires a panel of reference samples as auxiliary input, Polaris, once trained, does not rely on such panels (Fig. 1c), making it applicable to data across various species.

**Figure 1:**
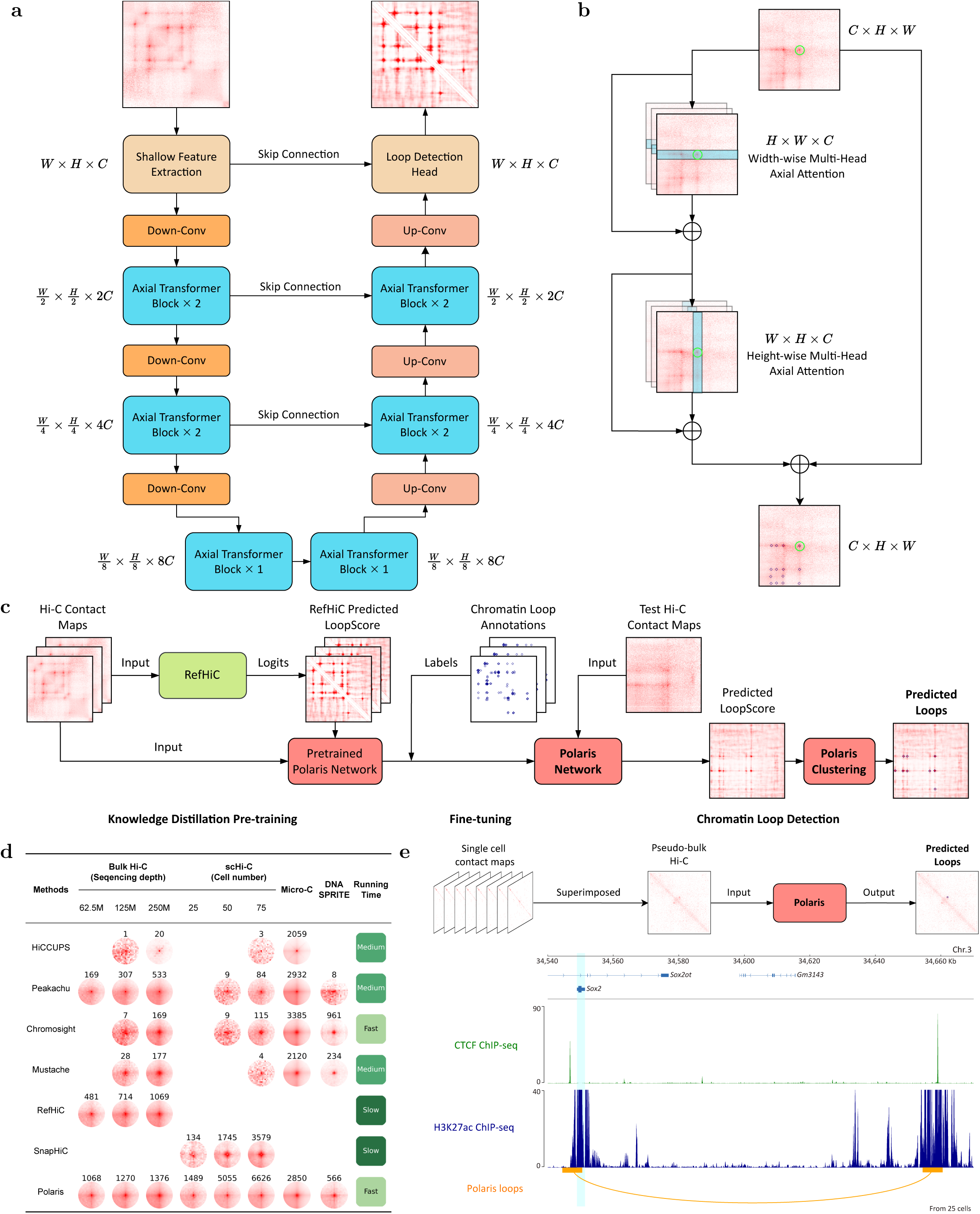
Polaris improves loop annotation in hold-out contact maps by leveraging axial attention and a large receptive field. **a,** Overview of the Polaris neural network for loop scoring. **b,** Schematic representation of the axial transformer block used in Polaris. **c,** Pretraining-finetuning paradigm for developing Polaris, along with steps for applying it to detect chromatin loops from chromosomal contact maps. **d,** Predicted loop number, aggregate peak analysis (APA), and running time comparison of Polaris with alternative tools for loop annotation from bulk and single-cell Hi-C, Micro-C, and DNA SPRITE datasets. For runtime efficiency, lighter green represent higher efficiency. **e,** Top: Application of Polaris for chromatin loop annotation from scHi-C data; Bottom: loop detected by Polaris from aggregated scHi-C data of 25 mES cells align with CTCF ChIP-seq and H3K27ac ChIP-seq signals around the Sox2 locus [41].

Polaris is a universal and efficient loop caller capable of detecting chromatin loops from contact maps generated by various assays, including Hi-C [5], scHi-C [7], Micro-C [6], and DNA SPRITE [40] (Fig. 1d). The Aggregate Peak Analysis (APA) plots indicate that loops detected by Polaris are not only more numerous but also exhibit comparable or even greater enrichment for chromosomal interactions compared to those identified by alternatives. Remarkably, when applied to pseudo-bulk scHi-C data, Polaris successfully identified a previously reported long-range interaction at the Sox2 locus [41] using data from only 25 cells (Fig. 1e).

In our study, we used human chromosomes 11 and 12 for validation, chromosomes 15–17 for testing, and the remaining autosomes for training. To prevent data leakage, all results presented are derived from test chromosomes. We performed analyses at 5 kb resolution unless indicated otherwise.

### Polaris detects loops from scHi-C data

We compared Polaris with SnapHiC [31], HiCCUPS [14], Peakachu [29], Chromosight [28], and Mustache [27], on scHi-C data (Fig. 1d, 2). Unlike SnapHiC, which directly processes multiple scHi-C contact maps, Polaris and other tools use pseudo-bulk data created by aggregating scHi-C datasets of the same cell type.

We applied them to analyze datasets consisting of contact maps derived from different numbers of single mouse embryonic stem cells (mESCs) [8]. As shown in Fig. 2a, Polaris is capable of detecting chromatin loops from as few as 25 cells, whereas other tools require substantially more data. As the number of cells increased, all tools identified more loops; however, only Polaris and Chromosight maintained a stable detection rate, indicating Polaris’s consistency across varying numbers of cells. We then compared loops predicted by each tool from datasets derived from varying numbers of cells against loops identified by CTCF ChIA-PET data to evaluate prediction accuracy (Fig. 2b, Supplementary Fig. 4). Across all settings, Polaris consistently predicted more CTCF ChIA-PET validated loops compared to alternatives, with the advantage being more pronounced in datasets with fewer cells. Moreover, loop anchors predicted by Polaris showed greater enrichment for CTCF binding sites compared to those predicted by other tools (Fig. 2c).

**Figure 2:**
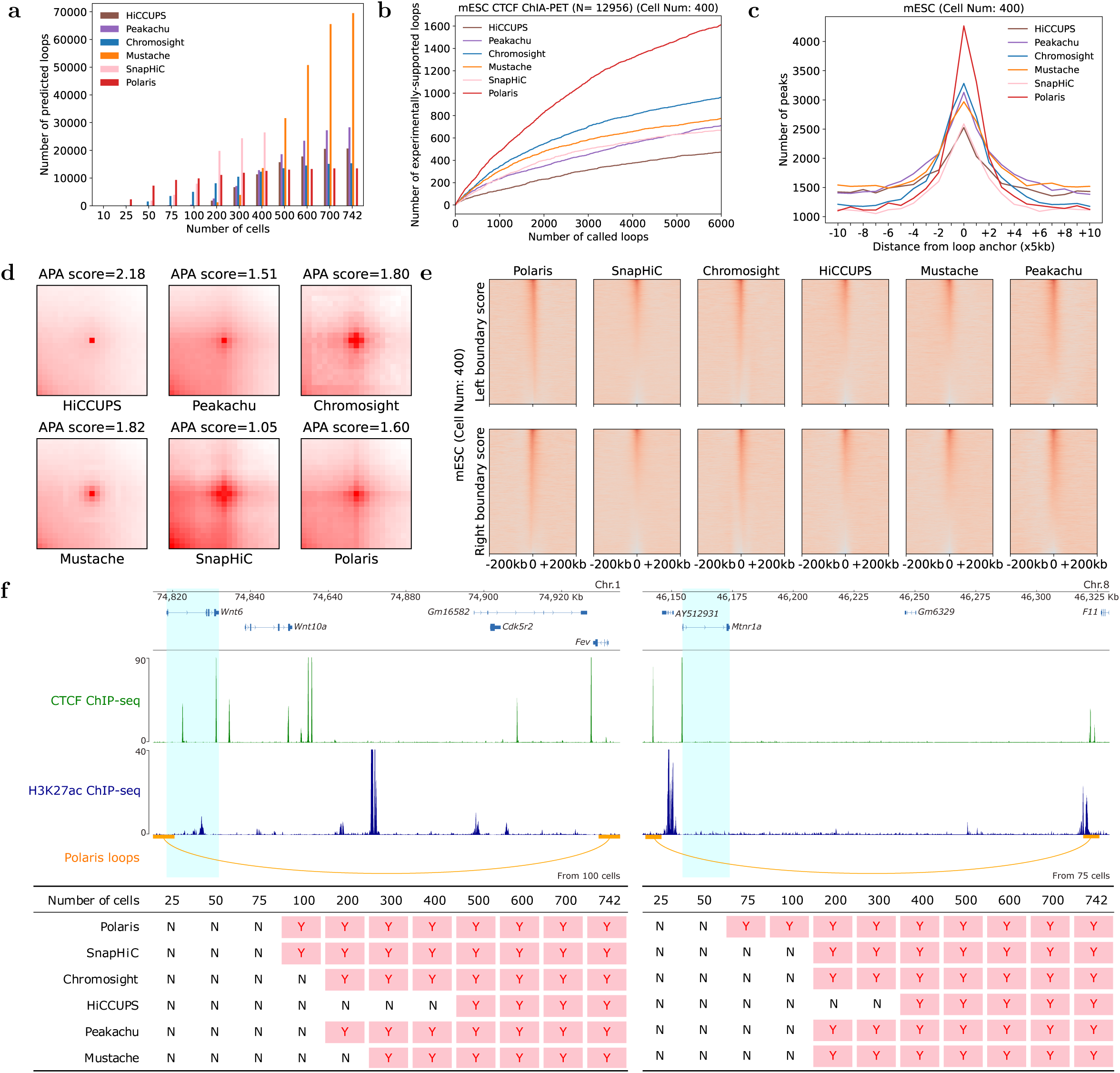
Performance of Polaris, SnapHiC, Chromosight, Peakachu, HiCCUPS, and Mustache on mESC scHi-C data. **a,** Number of loops detected by each tool from scHi-C data across varying numbers of cells. **b-e,** Comparison of loops annotated by each tool from scHi-C data derived from 400 mESCs. **b,** Number of CTCF ChIA-PET-validated loops among the top 6000 predictions made by Polaris and other tools. **c,** Occupancy of CTCF ChIP-seq signals around loop anchors annotated by each tool. **d,** Aggregate Peak Analysis (APA) for loops detected by each tool. **e,** Pile-up analysis of TAD scores around loop anchors predicted by each tool. **f,** Performance of each tool at the Wnt6 and Mtnr1a loci. Top: Loops around the Wnt6 (left) and Mtnr1a (right) genes, which have previously verified long-range interactions [42], identified from 100 and 75 mESCs by Polaris, with CTCF ChIP-seq and H3K27ac ChIP-seq as additional annotation tracks. Bottom: Comparison of tool performance in detecting such loops using various numbers of cells as input. ‘Y’ and ‘N’ indicate whether the previously verified interactions were identified or not.

The co-occurrence of loops, stripes, and TADs at the bulk level has been extensively investigated [14, 18]. However, these co-occurrences have not been thoroughly explored at the single-cell level. Here, we performed pile-up analyses of interactions around loops (Fig. 2d, Supplementary Figs. 5-6, Supplementary Note 2) and domain boundary scores around loop anchors (Fig. 2e, Supplementary Fig. 7, Supplementary Note 3). The APA results indicate that loops detected by Polaris are more enriched for chromosomal interactions and are often associated stripes. In contrast, loops detected by other tools were less associated with stripes and showed weaker blob-shaped patterns. The left and right domain boundary scores around corresponding anchors indicate that loops predicted by Polaris are more frequently located at TAD corners compared to those predicted by other tools.

To further assess the advantages of Polaris in situations with limited cell numbers, we conducted long-range interaction recovery analyses at the Sox2, Wnt6, and Mtnr1a loci, as previously identified in [41, 42]. As illustrated in Fig. 1e, 2f, and Supplementary Fig. 8, Polaris successfully detected these interactions from scHi-C data using as few as 25, 100, and 75 cells, respectively. In contrast, other tools required a larger number of cells to identify the same chromatin loops.

Taken together, Polaris stands out as a powerful tool for loop annotation from scHi-C data.

### Polaris detects loops from bulk Hi-C data

We evaluated the performance of Polaris against five loop callers: Chromosight [28], Peakachu [29], Mustache [27], HiCCUPS [14] and RefHiC [33] in bulk Hi-C data. We applied each tool to detect loops from downsampled Hi-C data (500M valid read pairs) derived from human GM12878 cells [14]. Loop predictions varied across tools, with Polaris identifying not only the highest number of loops but also the great number of unique predictions (Fig. 3a, Supplementary Figs. 9, 10a). Among unique predictions, those identified by Polaris were also more accurate compared to those by other tools (Supplementary Figs. 10b-e). Loops predicted by Polaris, Chromosight, Mustache, and RefHiC displayed more diffuse loop centers and visible stripes compared to those identified by other tools (Fig. 3b). Furthermore, the distances between loop anchors predicted by Polaris, RefHiC, and Mustache closely matched those of ChIA-PET/HiCHIP-supported loops, while other tools tended to favor shorter-range interactions (Fig. 3c). We then assessed the accuracy of each tool by comparing their predictions to those identified through loop-targeting experiment, allowing for a shift of up to one bin. To ensure a fair accuracy comparison, we selected the top 1700 predictions from each tool. As shown in Fig. 3d–f and Supplementary Fig. 11, Polaris detected 1271 CTCF-supported loops, 759 RAD21-supported loops, 557 SMC1-supported loops, and 213 H3K27ac-supported loops, comparable to predictions made by RefHiC. In contrast, other tools identified 17–53% fewer validated loops. Moreover, Fig. 3g shows that loop anchors predicted by Polaris and RefHiC were strongly enriched for CTCF binding sites. We also found that approximately 50% of the loops predicted by Polaris were enriched with convergent CTCF motifs, which is comparable to RefHiC and significantly outperforms other tools (Fig. 3h). A deeper analysis of the properties of loops predicted by each tool is presented in Supplementary Note 4 and Supplementary Figs. 12-13. To further study the performance on low coverage data, We benchmarked these tools on downsampled Hi-C contact maps, with valid read pairs ranging from 62.5M to 2,000M. As described in Supplementary Note 5 and illustrated in Supplementary Figs. 14-17, Polaris demonstrated consistent reliability in identifying loops across both low- and high-coverage Hi-C data. Last, we measured the running time of each tool in annotating Hi-C contact maps containing 4B, 1B, and 500M valid read pairs. Execution times varied from minutes to hours, with Chromosight and Polaris completing within 10 minutes—significantly faster than other tools (Fig. 3i).

**Figure 3:**
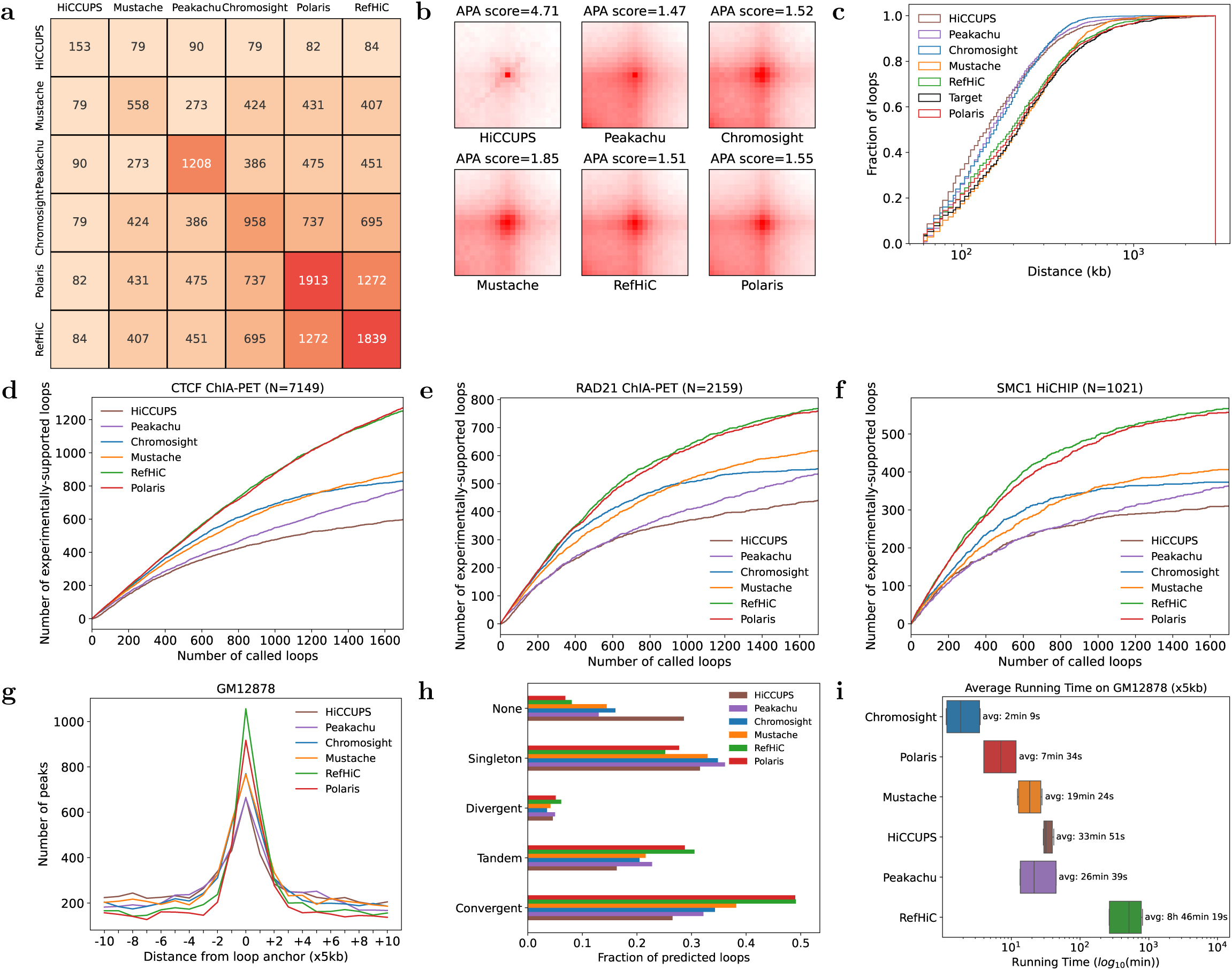
Performance comparison of Polaris, RefHiC, Chromosight, Peakachu, HiCCUPS, and Mustache on GM12878 Hi-C data (500M valid read pairs). **a,** Pairwise comparison of the number of loops detected by each tool. The value in each cell indicates the number of overlapping loops between the corresponding tools. **b,** Aggregate Peak Analysis (APA) for loops detected by each tool. **c,** Cumulative distributions of the distance between each pair of predicted loop anchors. The distribution of loops annotated by Polaris closely resembles that of loops supported by ChIA-PET/HiCHIP (target). **d-f,** Number of ChIA-PET/HiCHIP-supported loops among the top 1700 predictions made by Polaris and other tools, evaluated on test chromosomes 15-17. Comparisons are made with CTCF ChIA-PET (d), RAD21 ChIA-PET (e), and SMC1 HiCHIP (f) data. **g,** Occupancy of CTCF binding sites identified by ChIP-seq as a function of distance from predicted loop anchors. **h,** Orientation of CTCF motifs at predicted loops. **i,** Running time comparisons across different tools. The box plot shows execution times for each tool, measured five times under three settings (500M, 1B, and 2B valid read pairs), resulting in a total of 15 evaluations per tool.

Overall, these findings demonstrate Polaris’s superior performance compared to other tools in annotating loops from bulk Hi-C data.

### Polaris detects loops from DNA SPRITE and Micro-C data

To demonstrate the versatility of Polaris across assays, we applied it to GM12878 DNA SPRITE [43] and H1-hESC Micro-C [24] data. Polaris detected a substantial number of loops from both datasets, surpassing other loop callers except Chromosight (Fig. 4a). For DNA SPRITE, Polaris identified 692 loops, while Chromosight detected 1,184 loops, and other tools detected substantially fewer loops. For Micro-C, we observed a similar trend: Polaris detected 3746 loops, Chromosight identified 5,253 loops, whereas HiCCUPS and Peakachu detected about 3600 loops and Mustache identified 2,691 loops. For Micro-C data, which contains 2B valid read pairs, loops identified by all tools are enriched for chromosomal interactions and associated with stripes, as well as domain boundaries (Fig. 4b, Supplementary Figs. 18b, 19b). In contrast, for DNA SPRITE data, which only contains 50M valid read pairs, showed more variability (Fig. 4e, Supplementary Figs. 18a, 19a). Although tools like Mustache, and Polaris detected blob-shaped patterns, only Polaris’s predictions aligned with the presence of stripes and TAD corners.

**Figure 4:**
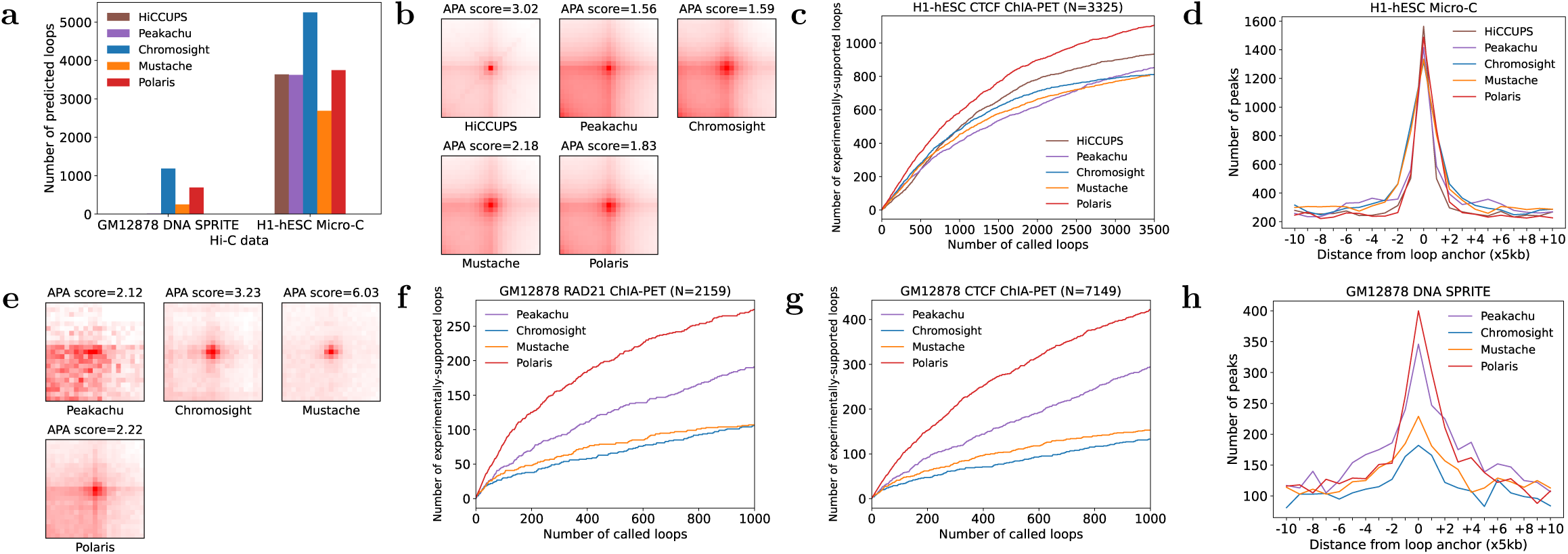
Detection of loops from GM12878 DNA SPRITE and H1-hESC Micro-C data. **a,** Number of loops detected by each tool from GM12878 DNA SPRITE and H1-hESC Micro-C datasets (test chromosomes 15–17 only) with recommended settings. **b,** Aggregate Peak Analysis (APA) for loops identified by each tool from H1-hESC Micro-C data. **c,** Number of CTCF ChIA-PET-supported loops among the top 3500 predictions made by Polaris and other tools. **d,** Occupancy of CTCF ChIP-seq signals in H1-hESCs as a function of distance from predicted loop anchors. **e,** Same as (b), but for loops annotated from GM12878 DNA SPRITE data. **f-g,** Number of RAD21 (f) and CTCF (g) ChIA-PET-supported loops among the top 1000 predictions made by Polaris and other tools from the DNA SPRITE data. **h,** Same as (d), but for loops annotated from GM12878 DNA SPRITE data.

We further evaluated the accuracy of loop predictions using ChIA-PET [44] and ChIP-Seq [45] data. For the Micro-C assay, among the top 3,500 loop predictions, Polaris detected more CTCF-supported loops compared to other methods, with 1,107 loops supported by orthogonal data (Fig. 4c). Furthermore, the occupancy of CTCF binding sites around predicted loop anchors in H1-hESC data showed that Polaris achieved high enrichment of CTCF ChIP-Seq signals at loop anchors, reinforcing its accuracy in detecting biologically meaningful chromatin loops (Fig. 4d). For the DNA SPRITE assay, we observed similar trends, with Polaris outperforming alternative tools in detecting loops validated by RAD21 and CTCF ChIA-PET and H3K27ac and SMC1 HiCHIP data (Fig. 4f-g, Supplementary Figs. 20-21), as well as showing anchors enriched for CTCF ChIP-Seq signals (Fig. 4h).

Overall, these results highlight Polaris’s robust performance in detecting chromatin loops from DNA SPRITE and Micro-C data, demonstrating its applicability to diverse assays.

### Polaris performs well across cell types and species

Although Polaris was trained by distilling knowledge from RefHiC [33], a reference panel-based method, it does not rely on reference panels during prediction. This enables Polaris to annotate loops purely from the study sample, making it applicable to data from various species, without being constrained by the availability of reference data.

We applied Polaris and other tools to Hi-C datasets from human K562, IMR90 [14], and cohesin-depleted HCT-116 [46] cell lines, as well as whole genome mESC [47] and zebrafish embryos (ZFE) [48]. We failed to apply RefHiC to analyze ZFE due to the absence of reference samples. Across all tools, we observed that higher-coverage data generally resulted in the recovery of more loops (Fig. 5a). On the K562 and IMR90 datasets, Peakachu, Chromosight, RefHiC, and Polaris identified similar numbers of loops, while HiCCUPS and Mustache detected 50–88% fewer loops. On mESC dataset (120M valid read pairs), Polaris, Peakachu, and RefHiC identified substantially more loops than other tools, highlighting their sensitivity to low-coverage data. For ZFE Hi-C data (234M valid read pairs), Polaris detected far more loops than other tools. We then assessed false positive rates by evaluating each tool’s performance on cohesin-depleted HCT-116 cells. Polaris, Peakachu, Mustache and RefHiC produced fewer than 50 loop predictions, indicating a low false positive rate.

**Figure 5:**
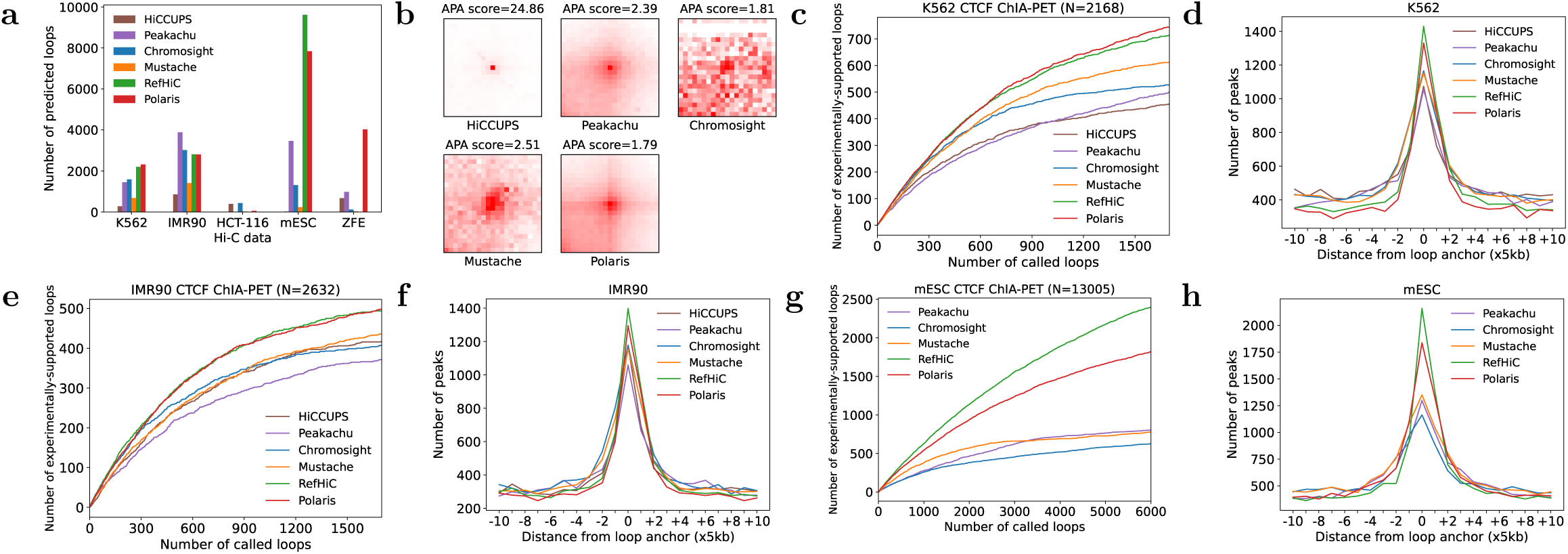
Detection of loops from Hi-C data in human K562, IMR90, cohesin-depleted HCT-116, mouse embryonic stem cells (ESCs), and zebrafish embryos (ZFE). **a,** Number of loops detected by each tool across various Hi-C datasets (for human data: test chromosomes 15–17; for mouse and zebrafish data: all autosomes) with recommended settings. Note that the cohesin-depleted HCT-116 dataset is expected to contain no loops. **b,** Aggregate Peak Analysis (APA) of loops detected by each tool in ZFE Hi-C data. **c,** Number of CTCF ChIA-PET-supported loops among the top 1700 predictions made by Polaris and other tools from K562 data. **d,** Occupancy of CTCF ChIP-seq signals in K562 cells as a function of distance from predicted loop anchors. **e-f,** Same as (c-d), but for loops detected in IMR90 Hi-C data. **g-h,** Same as (c-d), but for the top 6000 loop predictions from mESC Hi-C data.

To assess tool’s accuracy, we compared loop predictions with those identified through orthogonal experiments. For the K562 data (Fig. 5c, Supplementary Figs. 22d, 23a-b), Polaris demonstrated superior performance by identifying more loops supported by CTCF and RAD21 ChIA-PET. Additionally, loops predicted by Polaris are more enriched for CTCF binding sites than those predicted by alternatives (Fig. 5d). A similar pattern was observed in the IMR90 dataset (Fig. 5e-f, Supplementary Fig. 23c). For the mESC dataset (Fig. 5g-h, Supplementary Fig. 23d), although Polaris performed worse than RefHiC, it still outperformed others by a large margin, detecting twice as many CTCF-supported loops, with its loop anchors showing a stronger enrichment for CTCF binding sites. For the ZFE dataset, we employed APA and TAD score pile-up analyses to assess loop prediction accuracy (Fig. 5b, Supplementary Figs. 24d, 25d). Loops annotated by Polaris showed greater enrichment for chromosomal interactions, with anchors that were more frequently associated with domain boundaries. Same trends can be observed for other cells (Supplementary Figs. 22, 24, 25).

Overall, these findings highlight Polaris’s consistently superior performance across a range of cell types and species.

### Polaris performs well across different resolutions

To evaluate the impact of resolution on loop annotation, we applied Polaris and other tools to analyze GM12878 Hi-C data (500M valid read pairs) at additional 10kb and 25kb resolutions. As shown in Fig. 6a, the number of annotations varied across resolutions, with Polaris and Mustache showing a decrease in loop detection at lower resolutions. It indicates that Polaris and Mustache performed as expected, as higher resolutions provide finer details, allowing for the detection of more loops.

**Figure 6:**
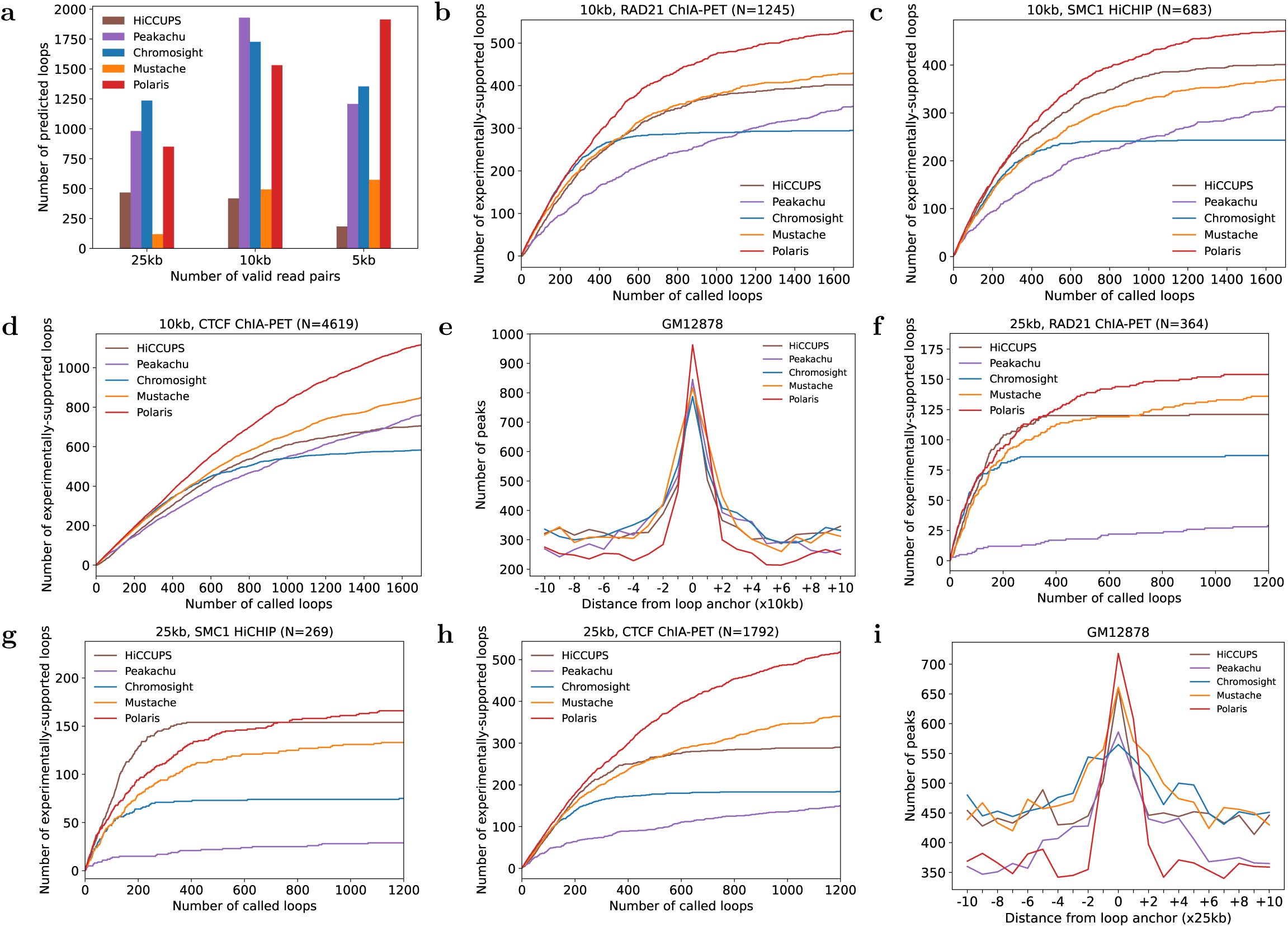
Comparison of tools on GM12878 Hi-C data at different resolutions (500M valid read pairs). **a,** Number of loops identified by each tool from Hi-C data across varying resolutions with recommended settings. **b-d,** Number of loops among the top 1700 predictions made by Polaris and other tools at 10kb resolution (test chromosomes 15-17 only) that are supported by orthogonal datasets: RAD21 ChIA-PET (b), SMC1 HiCHIP (c), and CTCF ChIA-PET (d). **e,** Occupancy of CTCF ChIP-seq signals as a function of distance from predicted loop anchors. **f-i,** Same as (b-e), but for loops annotated in Hi-C data at 25kb resolution.

We then evaluated tool accuracy using loops identified through orthogonal experiments. We selected top 1700, and top 1200 predictions from each tool’s outputs at 10kb and 25kb resolutions respectively. For the 10kb resolution, as shown in Fig. 6b-d and Supplementary Figs. 26c, 27, Polaris outperformed other methods in detecting loops validated by orthogonal data, similar to its performance at 5kb resolution. Additionally, Fig. 6e shows that loop anchors predicted by Polaris were more enriched for CTCF binding sites compared to those predicted by other tools. Similar trends were observed for predictions at 25kb resolution (Fig. 6f-i and Supplementary Fig. 26d, 27). We also observed Polaris consistently identified loops characterized by stripes and TAD boundaries (Supplementary Figs. 26, 28, 29). These findings underscore Polaris’s capability to detect chromatin loops across multiple resolutions.

## Discussion

Here, we introduce Polaris, a universal framework for chromatin loop annotation that demonstrates superior performance across a wide range of 3D genome mapping assays. Unlike existing tools, which are often optimized for bulk Hi-C data, Polaris is designed to analyze both single-cell and bulk datasets. It utilizes a novel neural network design combined with a pre-training and fine-tuning paradigm to maintain high accuracy across different resolutions and sequencing depths.

Polaris integrates axial attentions with a U-Net backbone, enabling it to detect chromatin loops by capturing multiscale features. This innovative design allows Polaris to effectively leverage axial features such as co-occurring loops, stripes, and TAD boundaries, thereby enhancing loop detection, particularly in sparse and single-cell datasets. Importantly, while Polaris integrates both local and global structural features (Supplementary Fig. 3), it remains capable of detecting loops that do not associate with long-range patterns (Supplementary Fig. 30). Additionally, our knowledge distillation-based pre-training addresses the long-standing challenge of limited high-quality training labels in this field.

Despite being trained primarily on bulk Hi-C data and fine-tuned on GM12878 Hi-C data at 5 kb resolution, Polaris demonstrates its effectiveness across diverse assays, sequencing coverages, and resolutions. It consistently outperforms other tools in detecting loops validated by orthogonal experiments. Furthermore, Polaris achieves this level of accuracy with remarkable computational efficiency, often running faster than alternative methods. This combination of accuracy and efficiency makes Polaris uniquely suitable for high-throughput 3D genomic analyses. While we observed slight improvements when fine-tuning Polaris on more specific data, such as very low coverage data and data at other resolutions, the gains were negligible, suggesting that retraining is generally unnecessary, especially given the impracticality of such data availability for emerging experimental technologies.

Although Polaris offers significant advancements, further improvements remain possible. Expanding training to encompass more diverse data sets could enhance its generalization ability. As Hi-C datasets grow in diversity, Polaris is well-positioned to harness this expanding data, paving the way for new insights into 3D genome organization.

Polaris is particularly well-suited for analyzing sparse scHi-C data and low-coverage datasets. We expect it to provide valuable insights into chromatin architecture at previously unattainable resolutions.

## Methods

### Polaris network architecture

The overall architecture of Polaris deep neural network is illustrated in Fig. 1a. It employs a U-shaped encoder-decoder backbone with skip connections to capture multi-scale features [38]. The encoder comprises a shallow feature extraction module, followed by three down-convolution blocks, each reducing the input width and height by 50%, with axial-attention [37] blocks following each down-convolution block. The decoder is symmetrically organized, which consists with three up-convolution blocks, each doubling its input width and height, with axial-attention blocks applied before each up-sampling operation, and a loop detection head to make the final prediction.

The shallow feature extraction module consists of two GELU activated convolution layers with a kernel size of three, designed to extract low level features from the input Hi-C matrix. It takes a Hi-C submatrix *M* ∈ ℝ^1^*^×H×W^* as input, and project it to embedding *H*_0_ ∈ ℝ^C×H×W^.

The three down-convolution and axial attention transformer pairs capture multi-scale long range features especially along axial directions. Specifically, each down-convolution reduces the width and height dimension by a factor of two, while doubling the number of channels. The axial-attention transformer block (Supplementary Fig. 1) consists of a LayerNorm (LN) layer, width-wise and height-wise multi-head axial attention modules, residual connections, and four GELU-activated convolution layers. We defined the axial-attention layer as follows:

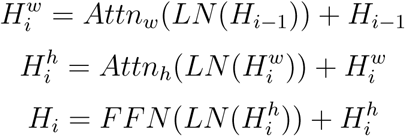

With 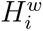, 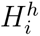, and *H_i_* represent the outputs of the horizontal, vertical, and axial attention in the *i^th^* axial attention block, respectively. The function *Attn*(*x*) refers to the *Q*, *K*, *V* attention operation as introduced by Vaswani et al. [49]. Similar to the encoder, the decoder is also built on axial-attention blocks. However, it incorporates up-convolution blocks, which consist of transpose convolution layers followed by GELU activation and a three-layer convolution module. These up-convolution blocks upsample the height and width of the learned features by a factor of two while reducing the feature dimension by half. The loop detection head consists of a single convolution layer with a kernel size of three, which predicts the loop probability (or score) for each pixel in the input Hi-C submatrix. To fuse multi-scale features from the encoder with the upsampled features in the decoder, skip connections are introduced between the corresponding layers of the encoder and decoder.

### Density-based clustering for loop detection from candidates

Polaris assigns loop score to each position in a Hi-C contact map using its deep learning module. To extract discrete loops from these scores, we use a clustering-based approach [50] to group loop candidates and identify representative loops. Specifically, pixels with loop scores above threshold t (*t* = 0.6) are treated as loop candidates. These candidates typically form clusters, from which we select one representative loop per cluster. Any isolated candidates (singletons) are likely false positives and are excluded from further analysis. For clustering, we use a density-based approach, similar to the method employed by RefHiC.

For each candidate (*i, j*), we first calculate its local density *ρ*(*i, j*) using a Gaussian kernel over the region within a Chebyshev distance of 5 bins. We then define *δ*(*i, j*) as the minimum distance to any candidate (*i^′^, j^′^*) with a higher local density. For the candidate with the highest density, we assign *δ*(*i, j*) = *δ*_max_, where *δ*_max_ is a constant. Candidates with *ρ*(*i, j*) above a defined threshold are marked as cluster centroids, with all other candidates assigned to their nearest centroid. Within each cluster, the candidate with the highest local density is predicted as a loop. Given the large number of loop candidates, calculating both *ρ*(*i, j*) and *δ*(*i, j*) can be computationally demanding. To address this, we employ a KD-tree data structure to efficiently accelerate the clustering process.

### Pre-training data and fine-tuning data

Polaris is designed to process matrices of dimensions 224 × 224. We pre-trained and fine-tuned Polaris on Observed/Expected (O/E) normalized Hi-C contact maps. Since chromatin loop anchors generally span distances of up to 3 Mb, we restricted our analysis to contact pairs within this range.

For pre-training, we curated a dataset of 25 human Hi-C contact maps at a 5 kb resolution, including seven GM12878 contact maps with varying numbers of valid read pairs (Supplementary Table 1). RefHiC was applied to these contact maps to produce logits for each pixel, which we used to compute soft labels for knowledge distillation. Soft labels were derived using the softmax function with a temperature parameter *T* :

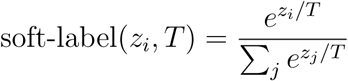

where *z_i_* is the logit for the *i*-th class, and *T* is the temperature parameter that controls the smoothness of the softmax output. In this loop detection task, which involves binary classification for each contact pair, we set *T* = 1, simplifying the soft label to:

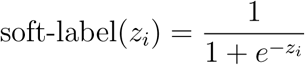

where *z_i_*is the logits predicted by RefHiC. We divided each Hi-C contact map and corresponding soft-label matrix into non-overlapping submatrices of size 224×224, discarding any submatrices with fewer than 10% non-zero entries. This process produced a total of 157, 456 samples for pre-training.

For fine-tuning, we utilized our established labeled loci from RefHiC to prepare training, validation, and test datasets. Briefly, we collected 74,855 positive targets, which represent loops identified across multiple experiments, including CTCF ChIA-PET, RAD21 ChIA-PET, SMC1 HiCHIP, and H3K27ac HiCHIP. We constructed 256,609 negative targets by drawing samples from non-loop loci in three categories: (i) 50,000 randomly sampled pairs with Chromosight scores below zero, matching the distance distribution of positive pairs; (ii) 10,000 randomly sampled pairs within a range of 1–3 Mb; and (iii) 196,609 loci identified as loops in RefHiC’s database but not in GM12878. The third group was specifically designed to train RefHiC, as RefHiC leverages additional reference samples, and we included this to ensure a fair comparison with RefHiC.

Following the same approach as RefHiC, we utilized a series of downsampled Hi-C datasets for GM12878 cells at 5 kb resolution as input. These targets were allocated into 224 × 224 submatrices, resulting in a total of 33,347 samples for training, validation, and testing.

### Pre-training via knowledge distillation

During pre-training, we used chromosomes 11 and 12 for validation, chromosomes 15–17 for testing, and the remaining autosomes for training. This dataset comprised 129,838 training samples, 14,800 validation samples, and 12,818 test samples. Training was conducted using a batch size of 480 and the AdamW optimizer [51] with a learning rate of 1e-4 over 1000 epochs on eight A800 GPUs. A warm-up strategy was employed for the first five epochs, progressively increasing the learning rate from zero to the target value, followed by cosine annealing to gradually reduce the learning rate to 1e-6 over the remaining epochs. To mitigate overfitting, dropout (ratio=0.1), and early stopping was applied. Furthermore, Exponential Moving Average (EMA) was utilized to smooth weight updates, enhancing model stability and generalization by reducing sensitivity to data noise. The network was pre-trained with the mean squared error.

### Supervised fine-tuning

We used the same criteria as in the knowledge distillation phase to split the fine-tuning dataset, which included 27,070 training samples, 3,232 validation samples, and 3,045 test samples. During fine-tuning, Polaris’s network initialized with weights obtained from knowledge distillation and was trained for 60 epochs using a batch size of 240 on eight A800 GPUs with the AdamW optimizer [51] with a learning rate of 1*e* − 6. To enhance model stability and generalization, we applied EMA, as in the pre-training phase. We followed RefHiC, and used focal loss −(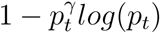) with *γ* = 2 to train our model.

### Hi-C data downsampling

To evaluate the impact of sequencing depth on Polaris’s ability to detect chromatin loops, we downsampled the Hi-C contact map of GM12878 cells. The combined Hi-C contact map for GM12878 cells was obtained from the 4DN Data Portal. Using the downsample function from FAN-C [52], we generated a series of downsampled datasets with valid read pairs ranging from 2,000M to 62.5M. Additionally, we used the zoomify function from Cooler [53] to create multi-resolution Hi-C contact maps at 5kb, 10kb, and 25kb resolutions.

### Loop detection from single cell Hi-C data sets with alternative tools

We downloaded single-cell Hi-C (scHi-C) data from mouse embryonic stem cells (mESC) to benchmark each tool’s performance in detecting loops from scHi-C data. Following the criteria set by SnapHiC, we included only Hi-C contact maps containing more than 150,000 valid read pairs for loop annotation, resulting in a dataset of contact maps for 742 cells. Similar to alternative tools, the scHi-C data at high resolution was too sparse to be analyzed effectively. Therefore, we created pseudo-bulk Hi-C contact maps by aggregating scHi-C data from cells of the same type.

To evaluate the performance of each tool, we generated a series of pseudo-bulk datasets containing contact maps from 10, 25, 50, 75, 100, 200, 300, 400, 500, 600, 700, and 742 cells, respectively. For tools such as HiCCUPS [14], Peakachu [29], Chromosight [28], and Mustache [27], which were originally developed for bulk Hi-C data with billions of raw reads, we performed parameter optimization on aggregated scHi-C data from 400 mESCs. For each tool, we identified a set of parameters that achieved the best performance on single-cell Hi-C data (Supplementary Note 6 and Supplementary Fig. 31). We then applied HiCCUPS, Peakachu, Chromosight, Mustache, and Polaris with optimized parameters to these pseudo-bulk contact maps. For SnapHiC [31], we directly applied the tool to scHi-C datasets containing the corresponding number of cells. We used 16 CPU threads and a single A800 GPU to run HiCCUPS, Peakachu, Chromosight, Mustache, and Polaris. For SnapHiC, due to its high computational demands, we allocated 50 CPU threads.

### Loop detection from bulk Hi-C data sets with alternative tools

We compared Polaris on bulk Hi-C datasets with five other tools: Chromosight [28], HiC-CUPS [14], Mustache [27], Peakachu [29], and RefHiC [33]. For tools like RefHiC and Peakachu, we trained them on the same labeled dataset used to train Polaris. We executed each tool with the recommended parameters, applying additional hyperparameter tuning as needed. In experiments where we controlled the number of predicted loops, we adjusted the FDR or loop score thresholds accordingly. All tools were run on the same server, utilizing 16 CPU threads and a single A800 GPU for inference.

## Data availability

All datasets utilized in this study are accessible to the public. Hi-C contact maps were obtained from 4DN Data Portal with the following accession code: 4DNFIXP4QG5B (GM12878), 4DNFI4DGNY7J (K562), 4DNFIJTOIGOI (IMR90), 4DNFILP99QJS (HCT-116), 4DNFIDA2WGV8 (mESC), 4DNFI8H9ECLI (ZFE), and 4DNFI6HDY7WZ (H1-hESC). ChIP-Seq data were collected from the ENCODE portal with the following accession code: ENCFF796WRU (GM12878 CTCF), ENCFF039JOT (GM12878 H3K27me3), ENCFF662DRZ (GM12878 RAD21), ENCFF887CRE (GM12878 SMC1), ENCFF508CKL (mESC CTCF), ENCFF203SRF (IMR90 CTCF), ENCFF119XFJ (K562 CTCF). The ChIA-PET data were obtained from ENCODE portal with accession code: ENCFF682YFU (IMR90 CTCF), ENCFF550QMW (mESC CTCF), ENCLB784HEF (GM12878 RAD21), ENCFF001THV (K562 CTCF). The RAD21 ChIA-PET for K562 were downloaded from the GEO repository with accession code GSM1436264.The H3K27ac HiCHIP data for GM12878 were obtained from ref. [54].The SMC1 HiCHIP data for GM12878 were obtained from ref. [55]. The CTCF ChIA-PET data for GM12878 were obtained from ref. [56]. H1-hESC Micro-C data and GM12878 DNA SPRITE data were obtained from 4DN Data Portal with accession code 4DNFI9GMP2J8 and 4DNFIUOOYQC3, respectively. The pre-processed scHi-C data of mES cells are publicly available at Tanay lab’s github repository.

## Code availability

Software is available at https://github.com/ai4nucleome/Polaris. Documentation is available at https://nucleome-polaris.readthedocs.io/en/latest. Source data and scripts required to reproduce analyses are available at https://zenodo.org/records/14294273.

## Acknowledgments

The authors thank Dr. Lujia Wang, Jiayong Zhou, Duohan Wu and Zhengjie Zhu for useful discussions in this project.

## Contributions

Y.Z., M.B., and Y.H. conceived the study. A.B. conceived single-cell data analysis. Y.Z. and M.B. supervised the project. Y.Z., and Y.H. designed models. Y.H. implemented models, performed data analysis. All authors wrote, read and approved the final paper.

